# Trophic niche partitioning in symbiotic marine invertebrates

**DOI:** 10.1101/2024.02.05.578332

**Authors:** Isis Guibert, Inga Elizabeth Conti-Jerpe, Leonard Pons, Kuselah Tayaban, Sherry Lyn Sayco, Patrick Cabaitan, Cecilia Conaco, David Michael Baker

## Abstract

Fierce competition for food and space underpins coral reefs’ biodiversity - supported by photosymbiotic foundational species. In contrast to other ecosystems, there is scant evidence that competition is mitigated by niche partitioning. Indeed, the dynamic evolutionary lineages of symbiotic partners and their syntrophy create layers of nutritional complexity that obfuscate patterns that structurn reef communities. As conspicuous members of Indo-Pacific reefs - giant clams co-occur with reef-building corals and similarly associate with algal symbionts. Using a common garden experiment, we analyzed stable isotope values from six giant clam species in the Philippines. These data, along with published data from ten sympatric corals, were used to calculate a novel metric - the Host Evaluation: Reliance on Symbionts (HERS) index - to assess variations in relative trophic strategies. Consistent with trophic niche partitioning – all species fell along an autotrophy-heterotrophy gradient with little overlap. We found a significant phylogenetic signal in clam HERS score, highlighting the role of selection in their nutritional ecology. We conclude that niche partitioning comes with tradeoffs, where predominantly autotrophic species showed higher growth rates but higher susceptibility to stress and consequently - greater conservation concern.

**Teaser:** Trophic niche partitioning plays a role in symbiotic marine invertebrate evolution with benefits and costs.

## Introduction

A fundamental question in ecology is what determines the number of organisms that can coexist in one environment(*1*). The concept of niche partitioning postulates that competition between species for scarce resources applies evolutionary pressure for novel adaptations – most famously in anatomical adaptations for acquiring nutrition ranging from animal dentition to plant roots (*2*). New traits that allow species to access uncontested resources reduce competition and provide an evolutionary advantage. Limits on how finely resources can be partitioned may therefore restrict the number and abundance of species in an environment(*3*).

The coevolution of inter-kingdom symbioses is a prevalent adaptive strategy that opens novel niche space to both partners(*4*, *5*). By coupling their metabolisms, symbiotic organisms increase the number of nutrient pools that can overcome nutritional deficiencies(*6*). Symbiosis allows holobionts (hosts and associated microorganisms) to thrive when resources are scarce, colonize new environments(*7*, *8*), and diversify(*9*). For example, plants colonized the terrestrial environment through coevolution with mycorrhizae, exchanging photosynthates for nitrogen, phosphorus, and water and thereby overcoming nutrient limitation(*7*). Novel niche space opened through symbiosis presents opportunities for additional partitioning, by varying the reliance on auto-trophy and heterotrophy contributing to the coexistence of species within ecosystems with varied environmental conditions(*10*). Nutritional symbioses are so successful that they form the foundation of several ecosystems including forests, deep sea vents, and coral reefs(*12*).

Coral reefs are home to unparalleled biodiversity, much of which is dependent on symbiosis to meet energetic requirements for growth and reproduction in a nutrient-poor environment(*13*). Oligotrophic habitats are where symbiosis thrives – as syntrophy confers a competitive edge over both seaweeds and suspension feeding invertebrates for limited space on the benthos. Many symbiotic marine invertebrates that dominate reefs are associated with dinoflagellate algae in the family Symbiodiniaceae – thus gaining access to photosynthates in exchange for metabolic wastes(*14*). Recent evidence suggests that within symbiotic taxa, dependence on syntrophic nutrients varies across host species(*9*, *15*). Corals, for example, fall along a continuum that ranges from a high dependence on autotrophy to complete heterotrophy(*15*, *16*). Trophic niche variation has been linked to colony morphology with branching and plating corals able to maximize their surface area available for symbiont photosynthesis while massive colonies are usually more heterotrophic with lower surface areas and larger polyps(*15*, *33*). Therefore, niche partitioning along an autotrophy-heterotrophy continuum can facilitate coexistence of species rich communities, achieving greatest diversity in places with dynamic sources of nutrition, particularly strong illumination and internal waves bringing plankton and inorganic nutrients (cite Wyatt Dongsha papers and there is a paleo paper that shows that mesotrophic conditions have greatest diversity).

However, this has only been explored in a few groups, and direct evidence of evolutionary niche partitioning within reef invertebrates remains limited(*9*).

Coexisting with corals, giant clams (subfamily Tridacninae) are symbiotic organisms consisting of 12 species divided in two genera. Tridacnids are ecosystem engineers -providing shelter, substrate, and food as well as benthic-pelagic coupling. Despite these important contributions, they remain understudied(*20*, *21*). Over the last decade, molecular and morphological phylogenetics have well supported the relationships within Tridacninae(*22–24*). Despite their relatedness, giant clams’ biogeography varies considerably, with a few species present broadly across the Indo-Pacific and others restricted to the center of the coral triangle(*25*). Depending on their location, giant clams can host one or more different genera of Symbiodiniaceae: *Symbiodinium*, *Cladocopium*, *Durusdinium* and/or *Gerakladium*(*26–28*). Unlike corals, giant clams establish extracellular symbiosis in their siphonal mantle and show less morphological variability with limited extension of their mantle. While aspects of the trophic capacity of seven giant clam species have been investigated using conventional techniques, these studies focused on a limited number of species (1 to 4) originating from different reefs(*29–31*) and used different methods, preventing cross-study comparisons. Trophic niche variation has been linked to size differences both within and between species(*29*, *31*), but previous studies have failed to compare the trophic strategies of giant clams of the same age or from the same environment, stressing the need for a comprehensive study.

The close physical and physiological integration between marine invertebrates and Symbiodiniaceae presents a challenge when assessing the relative contributions of each partner to the nutrition of the holobiont. Conventional methods for estimating the trophic niche of animals (gut content analysis and feeding experiments) are insufficient without also accounting for the translocation of nutrients from the photosynthetic symbiont (isotope tracer studies). The time-consuming nature of these approaches has restricted the number of taxa investigated. Recently, a relatively rapid method for assessing the trophic strategy of holobionts was developed using Stable Isotope Bayesian Ellipse in R (SIBER) analysis(*32*). This approach quantifies nutrient sharing and recycling within holobionts by measuring the overlap between host and symbiont “isotopic niches” as proxies for trophic niche. Previous applications of this technique have standardized overlap as a proportion of the area of the host niche, representing the contribution of translocated photosynthates to overall host nutrition(*33*, *34*). While this can adequately compare species when host and symbionts have similarly sized niches, it misses variation in the overlapping proportion of the symbiont ellipse that can occur when the symbiont ellipse is considerably larger or smaller than that of the host. Variation in the overlapping proportion of the symbiont niche is biologically relevant as an estimate of the impact of symbiont decline on host nutrition. Further, there is no consensus in the literature on what size niches should be used to compare groups (i.e. ellipses that encompass 40%, 75% or 95% of the bivariate isotope data)(*32*, *35*, *36*). We addressed these gaps by developing a novel index that incorporates multiple overlap metrics into one value indicative of host dependence on associated symbiont nutrition allowing for inter-species and inter-taxa comparison.

We employed this new metric to investigate trophic niche partitioning across six co- occurring species of giant clams and ten corals. All clams were sampled from a single lagoon in the Philippines whereas coral data was obtained from published studies(*33*, *37*). To gain a holistic picture of trophic niche partitioning in the giant clams, we characterized associated Symbiodiniaceae communities with next-generation amplicon sequencing to investigate functional differences. We further assessed whether nutrient sharing and other aspects of clam biology and ecology were linked to the phylogenetic relationships between clam hosts. We hypothesized that clams and corals would exhibit trophic resource partitioning along a heterotrophy-autotrophy spectrum explained by morphological differences. We also hypothesized that highly autotrophic clams would host unique or obligate symbiont associates. We expected functional traits including trophic strategy to show significant phylogenetic signals, supporting trophic niche partitioning as an evolutionary mechanism that drove divergence and speciation in Tridacninae. Finally, we predicted corals would exhibit a broader range of trophic strategies along the heterotrophy- autotrophy spectrum compared to giant clams, underpinning their extensive biogeography.

## Results

### Stable isotope analysis

We measured carbon and nitrogen stable isotope values of clam host tissue and associated Symbiodiniaceae in six co-occurring species from the Tabunan lagoon (Semirara, Philippines): *Hippopus hippopus*, *Hippopus porcellanus*, *Tridacna derasa*, *Tridacna gigas*, *Tridacna maxima*, and *Tridacna squamosa*. Extensive studies of symbiotic reef invertebrates including corals have shown that generally, d15N is an important predictor of trophic position while carbon is closely linked to energy sources (CITE). Host stable isotope values were generally higher than those of symbionts; mean δ^13^C ranged between -17.7±0.7 (*T. maxima*) and -12.3±0.9‰ (*T. derasa*) for clams and -18.9±0.9 (*T. maxima*) and -13.1±0.7‰ (*T. derasa*) for Symbiodiniaceae while mean δ^15^N ranged between 4.7±0.2 (*T. maxima*) and 5.2±0.3‰ (*H. hippopus*) for clams and 4.2±0.2 (*T. maxima*) and 4.9±0.1‰ (*T. gigas*) for symbionts (Supplementary Table 1). POM values averaged -18.9±1.7 for δ^13^C and 2.1±1.1 for δ^15^N (Supplementary 2).

To assess nutritional coupling between symbiotic partners, we first examined the difference between giant clam host and symbiont δ^13^C and δ^15^N (Δ^13^C and Δ^15^N) to investigate sharing of each nutrient individually(*10*, *33*). Both Δ^13^C and Δ^15^N varied significantly across species (Δ^13^C: Welch’s ANOVA, test statistic=10.397, P<0.0001; Δ^15^N: one-way ANOVA, F=16.3, P<0.0001; Supplementary Fig. 1). *T. gigas* Δ^13^C values were significantly lower than *H. hippopus*, *H. porcellanus*, and *T. squamosa* (Games-Howel Post-Hoc test, P<0.0001, P=0.0009, and P<0.0001, respectively) and *H. porcellanus* and *T. derasa* were significantly lower than *T. squamosa* (P=0.015 and P=0.001; Supplementary Fig. 1A). *T. gigas* Δ^15^N values were significantly lower than those of all other species (Tukey HSD, P<0.015), and *H. porcellanus* and *T. derasa* Δ^15^N values were significantly lower than *H. hippopus* and *T. squamosa* (P<0.005; Supplementary Fig. 1B).

Overall patterns of nutrient exchange were quantified by measuring the area of overlap of two differently sized Bayesian isotopic niches fit to bivariate δ^13^C and δ^15^N data: standard ellipse area set to encompass 40% each groups’ variation (SEAB; see Methods section for full definitions) and major ellipses area set to encompass 95% of each groups’ variation (MEAB)(*33*, *38*). Sample size of *T. maxima* (n=9) was lower than that of the other species (n=23 to n=28) simply because fewer individuals were available in the hatchery. While smaller sample size can affect ellipse area estimates, bootstrapped resampling approaches improve estimate reliability(*39*). We present the most robust analysis of our data given sampling constraints, however we encourage caution when interpreting the results of *T. maxima* due to its lower sample size.

Only one clam, *T. gigas*, exhibited overlap (0.29‰^2^) between host and symbiont SEABs, amounting to 24% of the host (SEABH) and 90% of the symbiont (SEABS) SEAB (Supplementary Table 2). All clam species, however, had overlap between host and symbiont MEABs, ranging from 0.41 (*H. porcellanus*) to 2.31‰^2^ (*T. gigas*; Supplementary Table 3). The overlap percentage of host MEABs (MEABH) ranged from 24 (*T. maxima*) to 49% (*H. porcellanus*) and exhibited less variation than that of the symbionts (MEABS), which ranged from 17 (*H. porcellanus*) to 98% (*T. gigas*). We also calculated the distance between host and symbiont ellipse centroids(*33*, *40*), which varied from 0.50 to 1.89‰ across species. Residual Permutation Procedures (RPPs)(*40*) demonstrated that host and symbiont groups occupied significantly different space on the isotope biplot for all species (P=0.001; Fig. 1B-F) except *T. gigas* (P=0.116; Fig. 1A).

**Figure 1.**
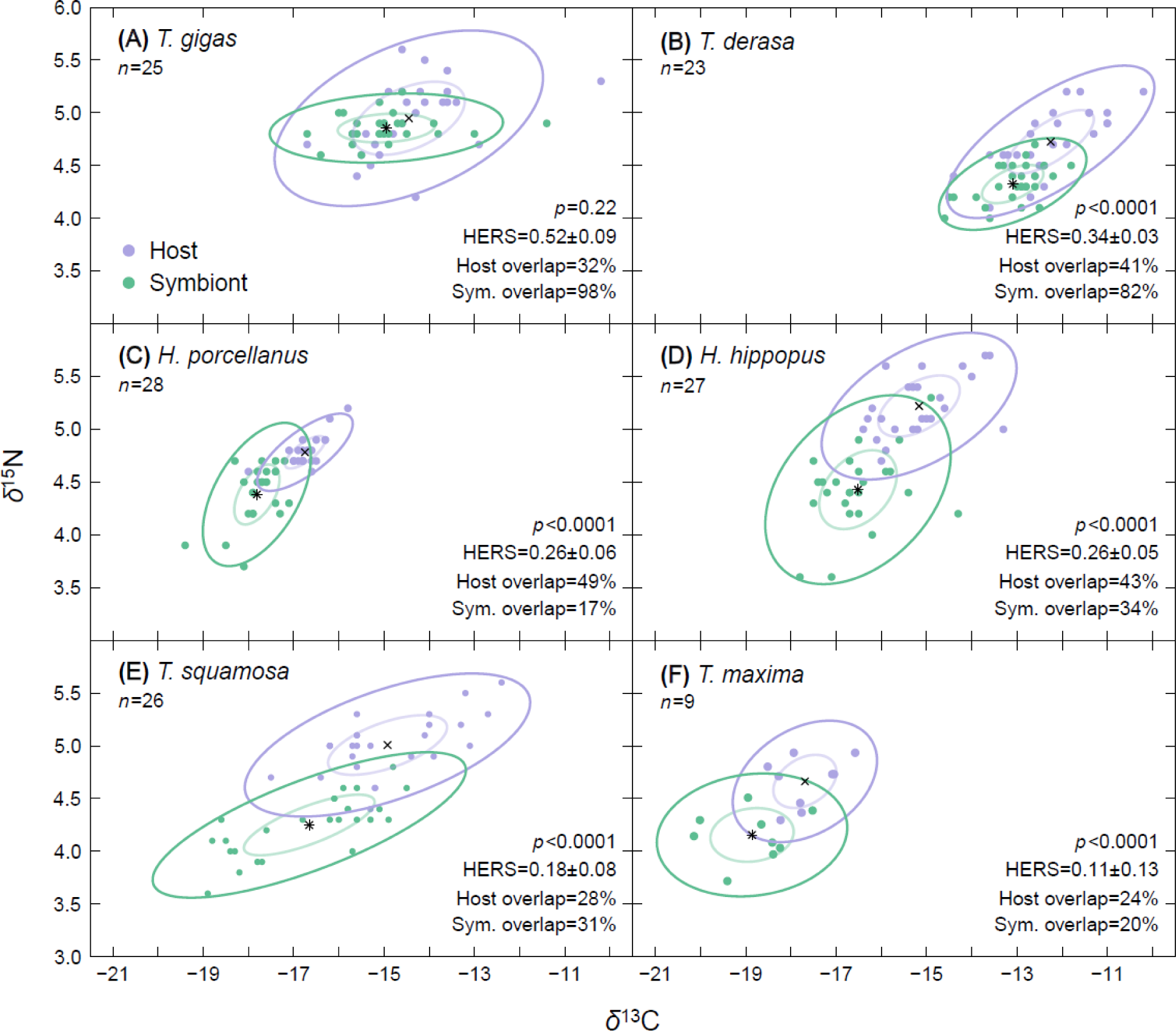
SIBER analysis of paired giant clam host (purple) and associated algal symbiont (green) stable isotope values. Lines represent ellipse areas corrected for sample size set to encompass 40% (standard ellipse area; SEA) and 95% (major ellipse area; MEA) of the variation of each group. Black symbols represent the centroids of host (*) and symbiont (x) ellipses. P-values<0.05 generated from residual permutation procedures (RPPs) indicate species where host and symbiont occupy distinct isotopic space. Upper and lower 89% credible intervals of each species HERS score indicate where they fall along a heterotrophy (0) - autotrophy (1) gradient. The percentage of overlap of host and symbiont MEAs were calculated using the mode of the last 100 posterior ellipses areas generated in a Bayesian analysis.

As a final metric quantifying nutrient sharing, we used a novel index (host evaluation: reliance on symbionts; ‘HERS’) that incorporated overlap values from both SEABs and MEABs to characterise the clams’ trophic strategy (see Methods section for full description). *Tridacna gigas* had a significantly higher HERS score (mean=0.52, 89% CI: 0.41-0.61) than all other clams and stood out as the only species with a mean score greater than 0.50 (Fig. 3). The species with the second highest HERS score, *T. derasa* (mean=0.35, 89% CI: 0.31-0.40), was significantly higher than the two species with the lowest HERS scores: *T. squamosa* (mean=0.18, 89% CI: 0.06-0.31) and *T. maxima* (mean=0.11, 89% CI: 0-0.30). The other two species (*H. hippopus*=0.26, 89% CI: 0.18-0.34; and *H. porcellanus*=0.26, CI: 0.18-0.37; Supplementary 2) had HERS values near the middle of the range across clams. Mean growth rates of clams species were found to correlate significantly with HERS scores (R^2^=0.94, P<0.01; Fig. 2). Using previously published data(*33*, *37*), we also calculated HERS scores for ten corals (*Acropora* spp., *Favites* spp., *Galaxea fascicularis*, *Gonipora* spp., *Pachyseris speciosa*, *Pavona* spp., *Platygyra* spp., *Pocillopora verrucosa*, *Porites* spp., and *Turbinaria* spp., Fig. 3). *Acropora* spp. had the highest HERS score close to 1 (mean=0.97) that was significantly greater (CI: 0.94-1.00) than all other corals. *Goniopora* spp., *Pachyseris speciosa*, and *Porites* spp. (mean=0.87-0.81) grouped together with scores significantly greater than corals with lower mean HERS scores (CI: 0.84-0.91, 0.81-.0.92, and 0.77-0.84, respectively). Corals with mid-range HERS scores (mean=0.34-0.62, *Pavona* spp., *Pocillopora verrucosa*, *Galaxea fascicularis*, *Favites* spp., *and Turbinaria* spp.) were significantly different from each other with three exceptions: *Pavona* spp. (CI: 0.57-0.67) and *P. verrucosa* (CI: 0.41- 0.61), *P. verrucosa* and *Galaxea* spp. (CI: 0.37-0.47), and *Favites* spp. (CI: 0.31-0.36) and *Turbinaria* spp. (CI: 0.30-0.37). The species with lowest score (*Platygyra* spp., mean=0.20) was also significantly different from all the other corals (CI: 0.17-0.23). The range of mean HERS scores across the ten coral species (0.20 to 0.97) was larger than that of the giant clams (0.11-0.52; Fig. 3; Supplementary 2). Four coral species (*Acropora* spp., *Goniopora* spp., *Pachyseris speciosa*, and *Porites* spp.) were significantly higher than all the clam species (corals’ CI: 0.77-1; clams’ CI: 0- 0.64). *Pavona* spp. and *Pocillopora verrucosa* were significantly greater than all the clams except *T. gigas* (corals’ CI: 0.57-0.61; clams’ CI: 0-0.40) and *Galaxea fascicularis* was significantly greater than *H. hippopus*, *T. squamosa* and *T. maxima* (corals’ CI: 0.37-0.47; clams’ CI: 0-0.34). *Favites* spp. was significantly different from *T. maxima* and *T. gigas* (corals’ CI: 0.31-0.36; clams’ CI: 0-0.30 and 0.41-0.61) while *Turbinaria* spp. was significantly lower than *T. gigas* (corals’ CI: 0.30-0.37; clams’ CI: 0.41-0.64) and *Platygyra* spp. was significantly lower than *T. gigas* and *T. derasa* (corals’ CI: 0.17-0.23; clams’ CI: 0.31-0.64).

**Figure 2.**
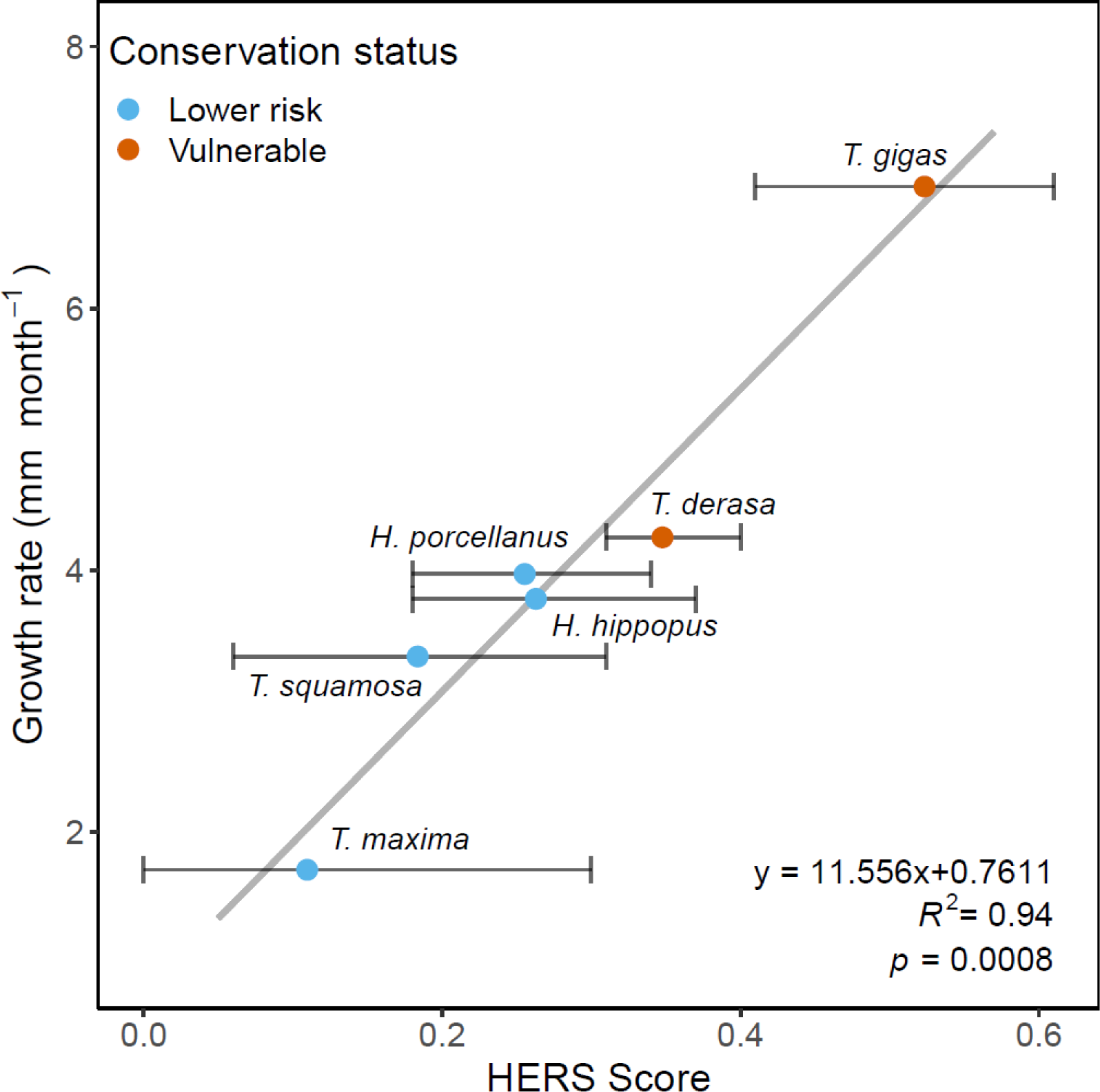
Significant correlation between the host evaluation: reliance on symbionts (HERS) scores and published mean growth rates of six giant clam species. Color indicates the International Union for Conservation of Nature (IUCN) conservation status of each species, which was either lower risk of extinction (blue) or vulnerable (orange)(*52*). 89% credible intervals are shown with dark gray error bars. Growth rates were obtained from Tan et al. (*23*).

**Figure 3:**
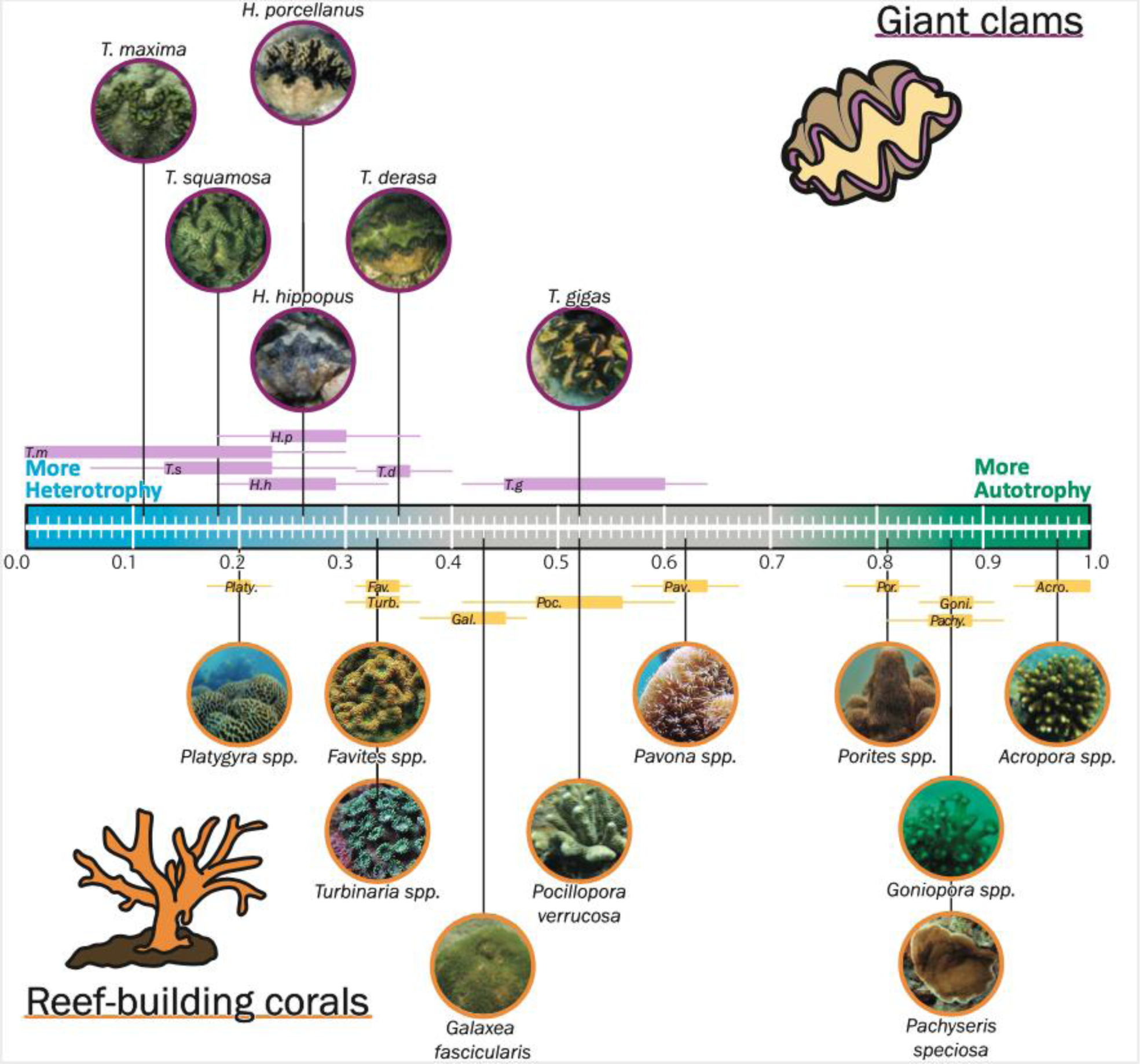
Host Evaluation: Reliance on Symbionts (HERS) scores of six giant clam species (topside of the scale; this study), three coral species, and seven coral genera (underside of the scale). Means HERS score is indicated by black lines connecting the species to the scale. Lower and upper quartiles of HERS score are indicated by horizontal boxes, and the credential interval low and high are displayed by lines. Coral stable isotope data used to calculate HERS values were obtained from Radice et al. (*37*) and Conti-Jerpe et al. (*33*). Drawings: Leonard Pons.

### Symbiodiniaceae diversity and community structure

We produced 1,890,847 sequence reads with an average of 105,047.10 (SD ±19,186) distributed over 18 giant clam samples which revealed 15 ITS2 type profiles belonging to three Symbiodiniaceae genera (Supplementary Fig. 2). Briefly, seven profiles belonged to the genus *Cladocopium* (78.21%), four were from genus *Durusdinium* (15.83%) and four were from *Symbiodinium* (5.96%). Only *H. hippopus* contained ITS2 type profiles representing all three genera while the other species hosted two genera (*Symbiodinium*-*Cladocopium* or *Cladocopium*- *Durusdinium*). Among the six species, *T. derasa* associated with highest number (seven) of predicted ITS2 type profiles, dominated by those from *Cladocopium* (C93a-C93e-C55a, 32.54%) and *Durusdinium* (D1-D4-D4c-D1c-D2, 31.39%) while *H. porcellanus* associated primarily with C93a-C93e-C55a (96.33%). Non-metric multidimensional scaling (NMDS) of ITS2 sequences and type profiles revealed distinct community structures with *T. maxima* and *T. squamosa* forming a distinct cluster (Supplementary Fig. 3).

### Phylogenetic signal analysis

We produced a Bayesian phylogenetic tree, based on 16 genetic sequences that corroborated evolutionary relationships between the six studied clam species supported in previous studies (Fig. 4)(*41*, *42*). Each node of the tree had a posterior probability greater than 0.89. The phylogenetic signal analyses showed significant patterns in four out of eight traits: mean growth rate (P=0.020; K=0.88), HERS score (P=0.027; K=1.04), maximum shell length (P=0.030; K=1.03) and depth (P=0.031; K=0.98).

**Figure 4.**
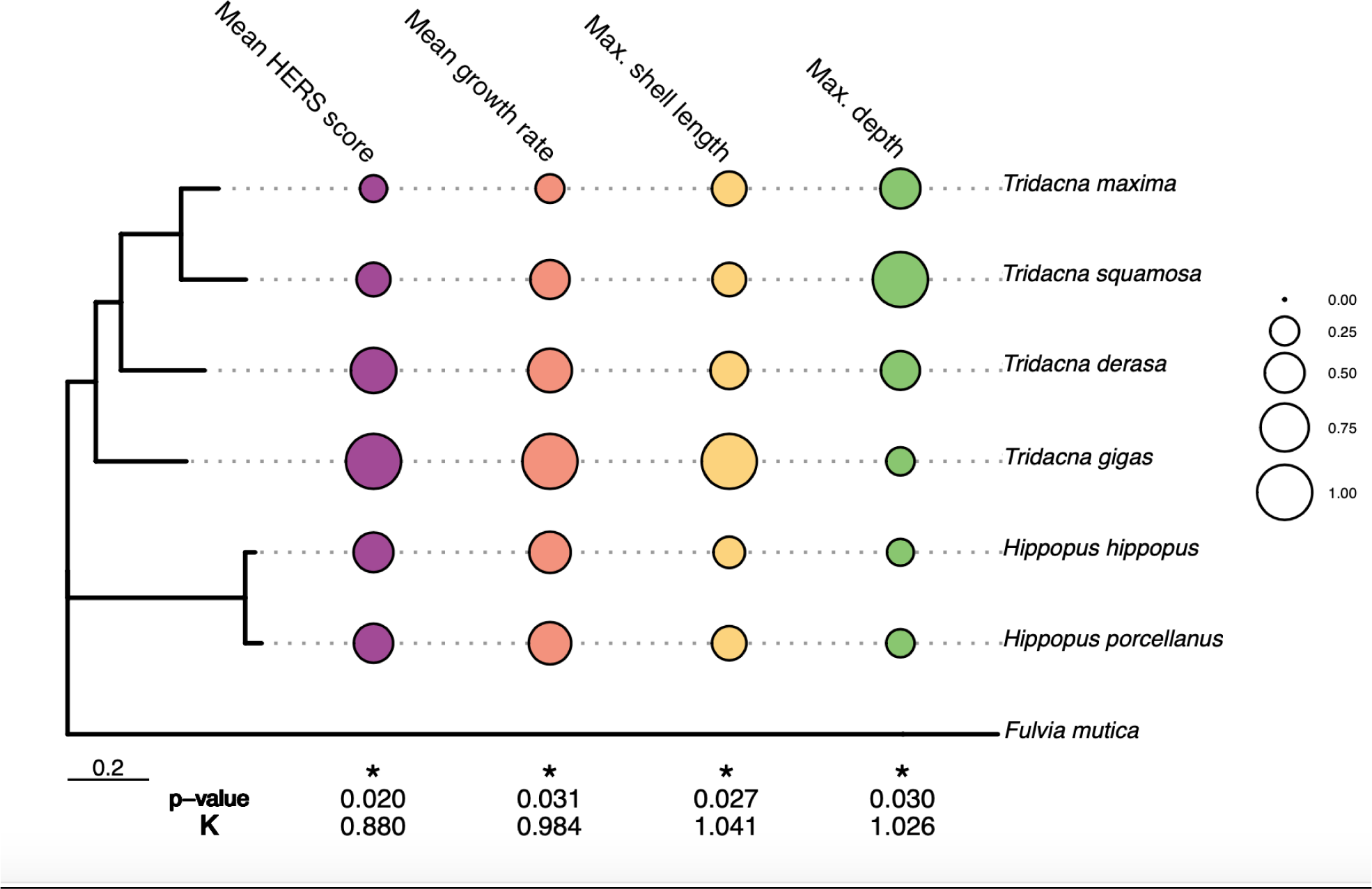
Significant (P<0.05) results of a phylogenetic signal analysis for six giant clam species. Traits with a significant phylogenetic signal include the mean HERS score, the mean growth rate, the maximum shell length and the maximum depth. The three traits aside from mean HERS score were obtained from Tan et al. (*23*). The size of each circle is proportional to the value used for each species in the analysis; the maximum value of each trait was fit to one and the minimum value to 0. The scale bar refers to the number of substitutions per site. *Fulvia mutica* is included as an outgroup species.

## Discussion

Giant clams are a remarkable group of marine invertebrates that are distributed across the Indo-Pacific and include the largest living bivalve on Earth, *T. gigas*. As large-bodied organisms that act as reef engineers by importing nutrients into benthic systems via suspension feeding, providing shelter or substrate through their shells, or acting as a food source for a wide range of predators, giant clams are crucial species in coral reef ecosystems(*21*). Yet they face many of the same anthropogenic threats as reef-building corals, including climate change, that affect their symbiosis and can lead to a rapid population decline(*20*). While corals have been extensively researched, giant clams are poorly studied despite their ubiquitous recognition in human culture and conservation concerns(*21*). As a taxon, they represent a unique case study to understand the role of nutritional symbioses in the evolution and biogeography of marine species.

The competitive exclusion principle postulates that species with overlapping niches compete(*2*), thus, subtle adaptations to avoid competition foster speciation and consequently, sympatry(*43*). Both clams and reef-building corals are considered mixotrophic organisms with the ability to obtain nutrition from both symbiont photosynthesis and suspension feeding(*44*) and both compete amongst themselves and with other photosynthetic and filter feeding taxa for food and space on oligotrophic reefs. We therefore hypothesized that variation in the relative contributions of auto- and heterotrophic dietary sources underpins the ecology and evolution of symbiotic marine invertebrates like giant clams, and helps explain their biology and biogeography across the Indo- Pacific. Further, we hypothesized that morphological traits predicate the trophic niche flexibility of symbiotic taxa with giant clams having less variation across trophic niches than modular corals.

By applying HERS to two broad taxonomic groups (Tridacninae and Scleractinia), we found evidence of trophic niche partitioning across mixotrophic marine invertebrates. Many species occupied statistically different regions of the HERS scale demonstrating variation in reliance on symbiont photosynthesis (Fig. 3). The lack of clear species groupings, particularly on the heterotrophic end of the scale, supports viewing mixotrophy as a continuum(*16*) rather than into category (i.e. predominant autotrophy, mixotrophy, and heterotrophy). While mutualisms with photosynthetic algae open additional energetic pathways to marine invertebrates, the evolution of this relationship across multiple taxa has resulted in fierce competition, providing a mechanism for the partitioning observed across clams and corals.

Giant clams did not extend as far along the heterotrophy-autotrophy spectrum as reef- building corals (Fig. 3), likely as a result of the high degree of morphological variability possible in colonial organisms(*45*). In corals, surface area to volume ratio is maximized through branching or plating colony morphology that is often seen in autotrophic species whereas more heterotrophic corals present massive colony shapes(*15*, *33*). In giant clams, symbionts are restricted to the ‘zooxanthellal tubular system’, which is exposed to light when the clam extends its mantle(*46*). Mantle size may therefore be an important morphological trait behind clam trophic variation, with larger mantles providing more surface area for photosynthesis. The inherent constraints on mantle size being restricted to the shell edge compared to the three dimensional flexibility of a coral colony could explain the relatively restricted range of clams along the trophic strategy spectrum. Giant clams are also able to filter large quantities of water (2000-28,000 Lh^-1^ of reef area) via siphonal inhalation(*21*), whereas corals must rely on ambient currents for particle capture. This likely explains the importance of heterotrophy to the trophic strategy of giant clams compared to corals.

Our results demonstrated that giant clams are likely constrained relative to corals on the autotrophy- heterotrophy spectrum, plausibly due to differences in functional morphology between both taxa.

Growth is an important tradeoff along the autotrophy-heterotrophy continuum. Corals that rely heavily on their Symbiodiniaceae often have faster growth rates(*47*), a pattern corroborated by our HERS results. *Acropora* spp., the most autotrophic coral with the highest HERS score, is fast- growing whereas more heterotrophic species such as *Platygyra* spp. grow slower(*48*). We observed a similar pattern in giant clams, where HERS scores correlated significantly with growth rates (P=0.0008; Fig. 2). Further, *T. gigas*, the most autotrophic clam that stood out with a HERS value >0.5, is able to achieve a body mass substantially larger than other clam species(*49*) of up to 200 kg(*50*) (Fig. 2 and 3). This size is supported by the photosynthesis of associated Symbiodiniaceae; growth rates of *T. gigas* can double at shallow compared to deep sites(*51*) and larger individuals cannot meet their energetic demands through heterotrophic feeding alone(*29*). Investigating trophic strategy across two taxa suggests that competition for both nutrients and space have played a role in the diversification of benthic reef organisms.

However, observations with both corals and clams suggest a tradeoff between autotrophy and survival in a changing world. *T. gigas* and *T. derasa*, the two most autotrophic clams, are also the only species in this study listed as Vulnerable on the IUCN Red List of Threatened species(*52*) (Fig. 2), suggesting higher sensitivity to global change. Similarly more autotrophic corals have been shown to be more susceptible to warming events(*33*). Indeed, in warming events around the world corals with branching morphology were the first to show signs of bleaching and exhibited the highest mortality and coral cover loss(*53*, *54*). These results indicate that while autotrophy likely supports high growth rates that confer fitness in space-limiting environments, marine invertebrates that rely heavily on this trophic strategy are less resilient to environmental perturbations, such as warming.

Using a unique “common garden” experiment, we demonstrated clear differences in trophic strategy across giant clams. Divergent strategies across species were supported by significant differences in HERS scores and other stable isotope metrics. Clams exhibited a range of HERS values; *T. gigas* was the most autotrophic and the only species with a value above 0.5 (0.52±0.09) while *T. maxima* was the most heterotrophic (0.11±0.13). The trend observed in HERS scores was corroborated by Δ^13^C and Δ^15^N values, which were both the lowest in *T. gigas*, supporting a high degree of autotrophy, and the highest in *T. squamosa*, *T. maxima*, and *H. hippopus*, indicating more dependence on heterotrophy. The distance between host and symbiont ellipse centroids was also shortest in *T. gigas*, which was the only species where host and symbiont groups did not occupy significantly distinct isotopic space. Given that the clams used in this work were all reared and collected from the same site characterized by an open, sandy bottom, the observed variation cannot be explained by environmental differences. These results provide evidence that giant clam species are not using identical trophic strategies and vary in their reliance on autotrophic and heterotrophic nutrient sources.

Across clam species, host nitrogen isotope values were offset from those of POM by the amount expected for a consumer (∼2.5‰)(*55*, *56*), suggesting that all species obtain some nutrition from filter feeding. Trophic differences across holobionts were instead driven by variation in the δ^15^N values of associated Symbiodiniaceae. In the most autotrophic species (*T. gigas*), the host and symbiont had near identical mean nitrogen values (Δ^15^N=0.1±0.3‰) whereas in more heterotrophic species (e.g. *T. maxima* and *T. squamosa*), symbionts had lower δ^15^N resulting in a larger difference from their hosts (Δ^15^N=0.5±0.3‰ and 0.7±0.3‰, respectively). This suggests that the mechanism for divergence of isotope-based metrics across clams was increased nutrient recycling between host and symbionts in more autotrophic species rather than a decrease in heterotrophy. Given the considerable active filtration capacity of clams(*21*, *57*), it is unsurprising that this nutrient source is used across species.

Recycling of nutrients within mixotrophic holobionts may eliminate isotope fractionation effects that could occur during nutrient translocation or excretion. It is unknown if fractionation occurs during nutrient exchange from Symbiodiniaceae to their invertebrate hosts. However, the similar δ^15^N and δ^13^C values of *T. gigas* hosts and symbionts suggest that either fractionation during translocation is minimal, or that nutrient recycling negates any fractionation effects. Similarly, depleted symbiont values relative to the host could result from the assimilation of isotopically light nitrogenous waste products the host preferentially excretes(*55*, *56*). Yet, if compounds produced with host waste are in turn shared with the host, these discrete isotope pools will be continually mixed, eliminating observable discrimination. While an understanding of the fractionation effects of the processes involved in syntrophic mutualisms would improve interpretation of our results, it is clear that host and symbiont partners with closer isotope values exhibit more nutrient sharing, recycling, or both than species with divergent values, regardless of the mechanism.

We posit that variation in giant clam nutritional strategy was underpinned by their evolutionary history. We detected a significant phylogenetic signal in mean HERS scores, supporting the hypothesis that trophic niche partitioning played a role in the divergence of species within Tridacninae(*58*). Previous work has found that the evolution of photosymbiosis in giant clams coincided with the expansion of the modern Indo-Pacific, implying that habitat was a key factor in the development of clam syntrophic relationships(*41*). Competition for nutrients in oligotrophic tropical seas applied selection pressure for the coupling of heterotrophic and autotrophic metabolisms, maximizing the potential avenues for meeting energetic requirements, while simultaneously opening an additional resource axis for partitioning that could reduce competition for scarce food(*59*). Previous studies found that up to six tridacnids can coexist on the same reef, however they occurred at different depths and substrate types, suggesting adaptation to microhabitats with varying light and/or particulate food availability(*60*). While multiple factors certainly contribute to the distribution of species, partitioning of resources, particularly those involved in both autotrophic and heterotrophic nutrient acquisition, contribute to the coexistence of clams and other sessile reef invertebrates such as corals, zoanthids, and sponges(*61*).

The ecological theory predicting increased selection for traits that lessen competition through niche differentiation is in accordance with our results. The phylogenetic analysis showed evidence of ecological divergence among the six species of giant clams based on functional traits, including growth rate and maximum shell length (a proxy for size)(*29*) (Fig. 3). More autotrophic clams, particularly *T. gigas*, exhibit faster growth and achieve larger body sizes. Growth rate and body size can be important adaptations and therefore play a role in evolution, for instance by increasing competitive abilities(*62*, *63*). Significant phylogenetic signal in the maximum depth was also revealed, with *T. maxima* and *T. squamosa* being able to colonize deeper environments (21.2m and 42m depth respectively) likely due to their predominantly heterotrophic strategy.

Symbiosis is another functional trait that plays a role in host ecology and evolution(*64*). Both environmental conditions(*19*) and host-intrinsic processes influence the composition of associated symbiont communities(*27*). Our results support clam host control over associated Symbiodiniaceae; six species harbored different genera and ITS2-type profiles despite inhabiting the same lagoon (Supplementary Fig. 2 and 3). Symbiont composition can also vary across a host’s geographic range. *T. maxima*, for example, associated with *Symbiodinium* in the Red Sea(*28*), *Cladocopium* and *Durusdinium* in the Philippines (this study; Fig. 4), and all three genera in French Polynesia(*26*). This flexibility coupled with low reliance on symbiont-derived nutrients could explain the wide geographic range of this species (Supplementary Fig. 5)(*25*). Regulation of associated symbiont communities may enable giant clams to meet their nutritional needs according to their environment without modification of the host’s physiology or morphology.

Although no studies have been conducted with clams, previous work with corals and anemones has demonstrated that the role of symbionts in host trophic dynamics varies according to Symbiodiniaceae genera(*67*). The two most heterotrophic species, *T. maxima* and *T. squamosa*, hosted the same genera, *Durusdinium* and *Cladocopium*, yet similar results were found for the predominantly autotrophic *T. derasa* species (Supplementary Fig. 2). While symbiont diversity did not appear to be linked to trophic niche variations between species, the ITS2-type profiles of *T. maxima* and *T. squamosa* were highly similar and distinct from all the other species (Supplementary Fig. 3). This indicates that while the Symbiodiniaceae likely play a role in giant clams trophic niches, future studies should focus on ITS2-type profiles.

Our study demonstrates that HERS is a useful tool for assessing the trophic strategy of mixotrophic holobionts and comparing their reliance on autotrophic and heterotrophic nutrients. HERS is a new, cohesive and robust metric that represents an integrated assessment of resource sharing, or the “strength” of the nutritional symbiosis. Palardy et al. (*16*) suggested that mixotrophic species should be considered on a continuum from 100% autotrophy to 100% heterotrophy, but until now existing metrics were unable to achieve this goal. While powerful, the multiple measurements provided by SIBER that reflect different facets of the trophic capacity of the holobiont (e.g. DEC, RPP and SEABH)(*33*), can be contradictory. For example, the distance between host and symbiont centroids in *T. gigas* (<1.0‰) indicated it is highly autotrophic whereas the overlapping proportion of host SEAB suggested moderate mixotrophy (24%)(*33*). The HERS scale addressed these discrepancies by incorporating both the proportion of host and proportion of symbiont niche overlapping that of the other partner, which encapsulates not only the host trophic regime but also the proportion of available photosynthate assimilated. This results in the most robust and biologically relevant quantifiable trophic indicator. Additionally, we included both the SEA (sensitive to autotrophy) and the MEA (sensitive to heterotrophy), in order to mitigate the bias inherent in using only one ellipse type as, seen in previous studies(*68–70*). Combining these measures into a single value raises the threshold for identifying holobionts that fall on either extreme end of the autotrophy-heterotrophy spectrum without assuming equivalent mixotrophy across syntrophic partnerships(*38*). These considerations resulted in a powerful metric that yielded a range of values consistent with previous findings and allowed for rapid and precise assessment of trophic niches (Fig. 3)(*33*, *37*).

Our research is key to understanding the trophodynamics of symbiotic marine invertebrates and therefore the relationship and energy flow of marine ecosystems. For the first time we demonstrated that trophic niche partitioning occurs within and across symbiotic taxa, providing an explanation for the co-occurrence of corals and giant clams on reefs. We posit that over evolutionary time, competition for scarce nutrients resulted in functional trait and trophic niche divergence, fostering coexistence. Our results also highlighted one potential tradeoff between reliance on autotrophy vs heterotrophy; more autotrophic giant clam and coral species grow faster yet are more susceptible to disturbance. Among the six species studied, the more autotrophic species (*T. gigas* and *T. derasa*) were listed as vulnerable on the IUCN Red List of Threatened Species demonstrating their sensitivity(*52*) and recent evidence shows that more autotrophic corals are more susceptible to bleaching(*33*). Therefore, while a commitment to symbiosis can enhance a holobionts competitive utilization of space - a clear advantage in nutrient- and space limited reef ecosystems, it is also an “Achilles’ heel” in an increasingly warm and eutrophic world. Trophic strategy may therefore be harnessed as an indicator of species of concern in conservation efforts(*71*). Giant clams and corals are also highly impacted by human activities including over-fishing and coastal development(*20*). Conservation of these ecosystem engineers, particularly susceptible species like *T. gigas*, *T. derasa* and *Acropora* spp., requires sustaining environmental factors that enhance autotrophic performance - mitigating seawater temperatures and reducing nutrients and sedimentation, specifically - in addition to curtailing over-harvesting.

## Material and Methods

### Sample collection

Six giant clam species (*Tridacna gigas*, *Tridacna derasa*, *Hippopus porcellanus*, *Hippopus hippopus*, *Tridacna maxima* and *Tridacna squamosa*) were spawned, matured and settled indoors at the Giant Clam Nursery of Semirara Marine Hatchery Laboratory before being transferred to sandy areas at 2-5 m depth in the adjacent Tabunan lagoon (Semirara Island, Philippines, 12°05’13.62” N, 121°20’45.66” E; Supplementary Fig. 5).

To explore niche partitioning across clams, we used a non-lethal sampling protocol in which one small (approximately 1 x 1 cm) mantle clip was harvested per clam. Mantle clippings were collected from 27 *T. gigas*, 28 *T. derasa*, 23 *H. porcellanus*, 25 *H. hippopus*, nine *T. maxima* and 26 *T. squamosa* juveniles and stored at -40 °C (Supplementary 3). Each mantle was clamped ∼0.5 cm from the edge using sterile hemostat forceps and snipped with sterile surgical scissors. Mean wet mass of clips varied from 0.095±0.035 g (*T. derasa*; ±SD) to 0.177±0.046 g (*H. porcellanus*) which was just enough for SIA analysis. Seawater (3 L) was collected and filtered to obtain particulate organic matter (POM; Supplementary 1 and 2). Frozen mantle clips and filters were shipped to the University of Hong Kong for subsequent analyses. Additional mantle clips were collected from three individuals of each giant clam species to characterise associated symbiont communities (Supplementary 3). Samples were stored in 5 mL screw-capped tubes filled with salt- saturated DMSO-EDTA buffer at 4°C until DNA extraction.

### Laboratory analyses

Mantle samples for SIA were lacerated with a razor blade, placed in 5 ml of Milli-Q water and sonicated (S-150D, BRANSON, United -States) for 30 s at 5 W to extract symbionts. The supernatant, containing the symbionts, was retained. This process was repeated to maximize symbiont yield. The combined symbiont fractions (10 ml) were centrifuged at 3,000 RCF for 10 min to rinse and pelletize the symbionts. The lacerated mantle was homogenized in 5 ml of Milli-Q water with a Tissue-Tearor (Ultra Turrax T25, IKA labortechnik staufen, Germany) and centrifuged at 2,000 RCF for 10 min to separate giant clam host tissue from any residual symbionts. The supernatant was collected and centrifuged again for 5 min at 2,200 RCF. The resulting pellet was discarded and the supernatant containing the clean host fraction was kept for analysis. Host and symbiont fractions were stored at -40°C overnight and then freeze-dried (Alpha 1-4 LCS-Plus, Christ, Germany). Dry clam host tissue (0.9±0.1 mg) and symbionts (0.7± 0.1mg) were weighed into tin capsules. POM from the Tabunan lagoon was scrapped off the filters and acidified in silver capsules using two applications of 5 μL of HPLC-grade 6N HCL. δ^13^C and δ^15^N stable isotope values were measured at the Stable Isotope Ratio Mass Spectrometry Laboratory at the University of Hong Kong with an environmental analyser (Eurovector EA3028, Italy) coupled to a stable isotope ratio mass spectrometer (Nu Instruments Perspective, UK) with accuracy better than 0.1‰ for both δ^13^C and δ^15^N. Vienna PeeDee belemnite (δ^13^C=0.011‰) and atmospheric nitrogen (δ^15^N=0.004‰) were used as standards for C and N, respectively.

Total genomic DNA from giant clams was isolated using a Qiagen DNeasy Plant Mini Kit (Qiagen, Germany) following the manufacturer’s protocol with minor adjustments. Briefly, ∼1 cm^2^ of mantle tissue was transferred to 1.5 mL microcentrifuge tubes, drained of excess preservation solution, and homogenized with a pestle before completing extractions according to the manufacturer’s instructions. DNA samples were quantified with a Nanodrop™ spectrophotometer (Thermofisher, USA) and sent to Macrogen Inc., South Korea for sequencing. Details on the library preparation, primers and PCR can be found in Supplementary 1.

### SIA statistical analysis

Host and symbiont isotopic niches were estimated using Stable Isotope Bayesian Ellipses in R (SIBER) analysis(*32*). A full description of the methods can be found in the supplemental materials. In brief, SIBER was used to generate Bayesian estimates of ellipse areas (EABs) fit to host and symbiont δ^13^C and δ^15^N values shown on an isotopic biplot. These ellipses represent the isotopic niche of each partner, which are proxies for trophic niche(*72*). The overlap of host and symbiont niches have previously been used to assess relative dependence on autotrophy vs heterotrophy(*33*, *34*). However, inconsistencies in the application of SIBER in the literature indicated the need to develop a novel metric that incorporates the most suitable outputs into one value that quantifies the relative trophic strategy of mixotrophic holobionts. Compiling multiple pertinent SIBER outputs integrates different nutritional elements of mixotrophic organisms, including aspects of both host and symbiont trophic niche. Our novel metric used two differently sized ellipses (encompassing 40% and 95% of the data respectively) that represent the minimum and maximum of biologically relevant previously published values(*32*, *35*), producing the most robust results and allowing for more fine-scale distinctions across species. Further, HERS includes overlap standardized not only as a proportion of the host niche but also as a proportion of symbiont niche. Both of these values have biological relevance; host niche overlap represents the relative contribution of the symbionts to host nutrition whereas the symbiont niche overlap represents the proportion of symbiont photosynthates that are passed to the host. The proportional overlap of host (EABH) and symbiont (EABS) 40% (standard ellipse area; SEAB) and 95% (major ellipse area; MEAB) Bayesian ellipses were combined into a novel index called Host Evaluation: Reliance on Symbionts (HERS) which estimates the relative nutritional importance of the symbionts to the host nutrition on a scale ranging from 0 to 1 (Equation 1).

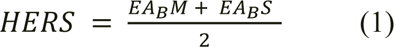

Where

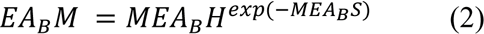

And

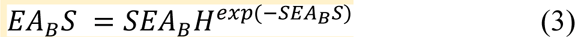

According to this index, HERS scores approaching 0 revealed limited nutrient sharing or recycling between host and symbionts, suggesting the host is more heterotrophic. In contrast, scores close to 1 suggested high exchange between the two partners, indicating the host is more autotrophic. Although high HERS scores by definition have high SEABH and MEABH results, the very top scores only represent holobionts that have, in addition, a higher proportion of SEABS and MEABS. This feature of the HERS score weighs the overlapping proportion of the host more than that of the symbiont (included as a negative exponent) resulting in a metric that is primarily informed by host niche overlap while still incorporating the proportion of symbiont niche overlap. Given that these niches reflect variation across individuals, complete overlap of the symbiont niche without complete overlap of the same sized host niche indicated variation in the divergence between host and symbiont isotope values.

Using stable isotope data we collected from the six clam species and published data from ten corals(*33*, *37*) (*Acropora spp.*, *Favites spp.*, *Goniopora spp.*, *Pavona spp.*, *Platygyra spp.*, *Porites spp.*, *Turbinaria spp.*, *Galaxea fascicularis*, *Pachyseris speciosa*, *Pocillopora verrucosa*) we ran 2000 independent SIBER analyses to generate 2000 HERS scores for each species (Supplementary 2). Boundaries of the 89% credible intervals (CI) were used to assess statistically significant differences between species(*73*).

We validated our HERS results for clams by calculating additional metrics previously used to evaluate the trophic strategy of mixotrophic organisms. To assess the sharing of carbon and nitrogen independently, we calculated Δ^13^C and Δ^15^N by subtracting the isotope value of the symbiont from that of its paired host(*10*, *33*). The distributions of Δ^13^C and Δ^15^N were assessed visually with Q-Q plots and statistically with Shapiro-Wilk tests, while scedasticity was evaluated with either a Bartlett’s test if data were normal or a Levene’s test if data were not normally distributed. Statistical differences between the Δ^13^C and Δ^15^N of different species were determined using one-way ANOVAs for normally distributed and homoscedastic data, or Welch’s ANOVAs for non-normal and heteroscedastic data. If significant differences between species were detected, pairwise comparisons were evaluated using either Tukey HSD tests or Games-Howell Post-hoc tests (Supplementary 1). The distance between host and symbiont ellipse centroids was also calculated with lower values indicative of more autotrophy and higher values indicative of more heterotrophy(*33*). Finally, significant differences in the relative placement of host and symbiont isotopic niches were determined using residual permutation procedures (RPPs)(*40*). We examined whether clam HERS scores were correlated with published growth rates(*74–76*) using a linear regression (generalized linear model). All isotope analyses were conducted in R version 4.2.1(*77*).

### ITS2 sequence analysis

Sequence analysis was conducted in the SymPortal framework (symportal.org)(*78*) to predict ITS2-type profiles as proxies for putative Symbiodiniaceae genotypes. Raw paired reads were submitted to the framework and subjected to quality control (QC) using mothur v.1.39.5(*79*).

Reads were screened for Symbiodiniaceae sequences within the range of 184–310 bp and algorithmically searched in a genus-separated manner using the BLAST+ suite of executables(*80*) and Minimum Entropy Decomposition (MED)(*81*). Post-QC, ITS2 sequences were loaded to both local and remote SymPortal databases to identify specific sets of ITS2 sequences re-occurring in multiple samples called defining intragenomic variants (DIVs). The presence and abundance of these DIVs were used to define the ITS2-type profiles. All sequences generated here were deposited at the National Center for Biotechnology Information (NCBI) under the BioProject ID PRJNA749183. To visualize Symbiodiniaceae community structures, non-metric multidimensional scaling (NMDS) based on Bray-Curtis dissimilarity measures from abundance data was plotted (Supplementary 1).

### Phylogenetic Analysis

Results were paired with a giant clam phylogeny to produce a phylogenetic signal analysis and investigate evolutionary patterns of ecological divergence among species(*9*, *14*). Mitochondrial and ribosomal sequences of the six studied giant clams and a cardiid outgroup species (*Fulvia mutica*) were obtained from Tan et al. (*23*). Model substitution for each of the 16 gene partitions were selected with the Bayesian information criterion (BIC) using jModelTest v2.1.10 (Darriba et al. 2012). A dated phylogenetic tree was constructed with MrBayes version 3.2.7a(*82*) from four independent Markov Chain Monte Carlo (MCMC) analyses, each containing four chains of two million cycles sampled every 100 cycles and trimmed from the first 25%. The phylogenetic tree was drawn from the resulting posterior distribution(*83*).

The multiPhylosignal function from the R package picante, was used to detect significant phylogenetic patterns in eight different ecological traits (Supplementary 5), using a bootstrapping technique with 720 (6!) replacements(*84*). To mitigate uncertainty produced by the permutation nature of this analysis, each final trait result was extracted from the mean of 1000 independent multiPhylosignal analyses (Supplementary material). Of the eight traits, three were obtained from the literature (mean growth rate, maximum shell length and maximum depth)(*23*), one (province’s occurrence) was determined by matching published occurrences of giant clam species(*25*) to established Marine Ecoregions of the World(*85*) resulting in the number of ecoregions each clam inhabits (Supplementary material 5), and four were determined in this study (DEC, mean δ^15^N, mean δ^13^C, and HERS score). K values greater or equal to 1 demonstrated a close fit to the phylogenetic relationship between species while values close to 0 indicated a pattern different from the phylogenetic tree(*86*, *87*).

## Acknowledgments

We deeply thank Dr. R. Estrellada, Director of the Semirara Marine Hatchery Laboratory (SMHL), as well as the staff of SMHL for hosting Dr. I. Guibert and for field assistance; J. I. P. Baquiran for logistics in the field; O. So for helping with all necessary orders; A. Y. T. Chan for technical assistance at the University of Hong Kong; Dr. O. Habimana for lending materials; Dr. S. McIlroy for valuable discussion; and M-J. Ho, Y-D. Pei and Dr. P. Thompson for granting us the use of their beautiful pictures.

## Funding

This study was funded by the University of Hong Kong Division of Ecology and Biodiversity PDF Research Award, by the Research Grants Council Collaborative Research Fund (17108620), by the Environment and Conservation Fund (ECF-67/2016 and CRF7G_C7013-19G) and by the Department of Science and Technology Philippine Council for Agriculture, Aquatic and Natural Resources Research and Development (DOST-PCAARRD; QMSR-MRRD-MEC-295-1449, 314-1542 and 314-1545).

## Author contributions

I.G. designed the study. I.G and D.M.B secured funding for the study. I.G, S.L.S, P.C. and C.C. carried out field sampling and helped with logistics. L.P., I.G., I.C.J and K. T. performed the laboratory work. L.P., I.C.J., K.T. and I.G. performed the analysis. I.G and I.C.J wrote the first draft of the manuscript and all authors contributed to the final manuscript.

## Competing interests

The authors declare no competing interest

## Data and materials availability

All sequences generated for this study were deposited at the National Center for Biotechnology Information (NCBI) under the BioProject ID PRJNA749183. All other data needed to evaluate the conclusions in the paper are present in the paper and/or the Supplementary Materials. Additional data related to this paper may be requested from the authors.

## Supplementary Materials

Supplementary 1 - Results, discussion and methods (Table 1-5, Figure 1-4)

Supplementary 2 - Stable isotopes data and HERS scores

Supplementary 3 - Description of the giant clam samples

Supplementary 4 - Most abundant ITS2 sequences

Supplementary 5 - Traits for phylogenetic analysis

## Notes

### Competing Interest Statement

The authors have declared no competing interest.

