## Supplementary for "Trophic niche partitioning in symbiotic marine invertebrates"

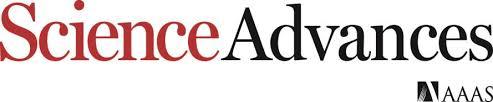

Supplementary Materials for

**Evolution of trophic niche partitioning in symbiotic marine invertebrates**

Isis Guibert*†, Inga Elizabeth Conti-Jerpe†, Leonard Pons, Kuselah Tayaban, Sherry Lyn Sayco, Patrick Cabaitan, Cecilia Conaco, David Michael Baker*.

†co-first authors

**This PDF file includes:**

Supplementary Text

Figs. S1 to S4

Tables S1 to S5

References (1 to 11)

**Summary:**

**Results**

Stable isotope analysis

Symbiodiniaceae diversity and community structure

**Discussion**

**Materials and Methods**

Sample collection

Habitat assessment

Laboratory analysis

SIA statistical analysis

**References**

**Other Supplementary Materials for this manuscript include the following:**

Data S2 to S5

**Supplementary Text**

**Results**

Stable isotope analysis

**Supplementary Table 1.** Summary statistics of stable isotope analysis of paired clam host and algal symbiont samples. Mean (±SD) carbon to nitrogen ratio (C:N), carbon (*δ*^13^C) and nitrogen (*δ*^15^N) stable isotope values, and difference between host and symbiont isotope values (Δ^13^C and Δ^15^N) of six giant clam species collected from Cover Bay, Semirara Island, Philippines.

| **Species** | **n** | **C:N** | | ***δ*^13^C (‰)** | | ***δ*^15^N (‰)** | | **Δ^13^C (‰)** | **Δ^15^N (‰)** |
| --- | --- | --- | --- | --- | --- | --- | --- | --- | --- |
|  |  | **Host** | **Symb** | **Host** | **Sym** | **Host** | **Sym** |  |  |
| *T. gigas* | 27 | 5.7±1.3 | 5.7±1.3 | -14.4±1.2 | -15.0±1.1 | 4.9±0.3 | 4.9±0.1 | 0.5±0.5 | 0.1±0.3 |
| *T. derasa* | 28 | 5.6±1.4 | 6.0±0.7 | -12.3±0.9 | -13.1±0.7 | 4.7±0.3 | 4.3±0.2 | 0.8±0.7 | 0.4±0.3 |
| *H. porcellanus* | 23 | 6.0±0.7 | 5.7±0.3 | -16.7±0.4 | -17.8±0.5 | 4.8±0.1 | 4.4±0.3 | 1.1±0.4 | 0.4±0.3 |
| *T. maxima* | 9 | 7.0±2.2 | 6.4±1.5 | -17.7±0.7 | -18.9±0.9 | 4.7±0.2 | 4.2±0.2 | 1.2±0.9 | 0.5±0.3 |
| *H. hippopus* | 25 | 5.5±1.2 | 5.5±0.5 | -15.2±0.9 | -16.5±0.8 | 5.2±0.3 | 4.4±0.4 | 1.3±0.6 | 0.8±0.4 |
| *T. squamosa* | 26 | 4.8±1.7 | 6.1±0.6 | -14.9±1.3 | -16.7±1.4 | 5.0±0.3 | 4.2±0.3 | 1.7±0.9 | 0.7±0.3 |

**Supplementary Table 2.** Standard ellipses areas and overlap metrics of six giant clam hosts and their associated algal symbionts in isotopic space. Stable Isotope Bayesian Ellipses in R (SIBER) analysis was used to fit standard ellipses area corrected for sample size (SEA*_C_*) that captured 40% of the variation of the data to host (Host SEA*_C_*) and symbiont (Symbiont SEA*_C_*) isotope values. The mode of the last 100 posterior ellipses from a Bayesian generated distribution was used as to compute the Bayesian standard ellipse area of the host group (Host SEA*_B_*), the symbiont group (Symbiont SEA*_B_*), and the area of overlap between the two. The area of overlap of host and symbiont was expressed as a proportion of Host SEA*_B_* (SEA*_B_*H) and Symbiont SEA*_B_* (SEA*_B_*S).

| **Species** | **Host SEA_C_ (‰^2^)** | **Host SEA_B_ (‰^2^)** | **Symbiont SEA_C_ (‰^2^)** | **Symbiont SEA_B_ (‰^2^)** | **SEA*_C_* area of overlap (‰^2^)** | **EA*_B_* area of overlap (‰^2^)** | **SEA_B_H** | **SEA_B_S** |
| --- | --- | --- | --- | --- | --- | --- | --- | --- |
| *T. gigas* | 1.19 | 1.13 | 0.43 | 0.41 | 0.40 | 0.29 | 0.24 | 0.90 |
| *T. derasa* | 0.60 | 0.59 | 0.34 | 0.32 | 0.10 | 0.00 | 0.00 | 0.00 |
| *H. porcellanus* | 0.14 | 0.14 | 0.39 | 0.37 | 0.00 | 0.00 | 0.00 | 0.00 |
| *T. maxima* | 0.51 | 0.41 | 0.73 | 0.60 | 0.00 | 0.00 | 0.00 | 0.00 |
| *H. hippopus* | 0.70 | 0.65 | 0.93 | 0.88 | 0.00 | 0.00 | 0.00 | 0.00 |
| *T. squamosa* | 0.95 | 0.87 | 0.89 | 0.86 | 0.00 | 0.00 | 0.00 | 0.00 |

**Supplementary Table 3.** Comparison of the placement and overlap in isotopic space of six giant clam hosts and their associated algal symbionts. Host and symbiont placements were assessed by measuring the distance between the ellipse centroids of the two groups (DEC) and running a residual permutation procedure (RPP). Host and symbionts occupied distinct space on the isotopic biplot when p<0.05. Overlap between the two groups was assessed using a Bayesian analysis to generate a posterior distribution of ellipses fit to encompass 95% of the variation of each group. The mode of the last 100 posterior ellipses and their overlap was used as the Bayesian ellipse area of the host group (Host MEA*_B_*), the symbiont group (Symbiont MEA*_B_*), and the area of overlap between the two. The proportion of host and symbiont MEA*_B_* overlapping that of the other was also calculated (respectively, MEA*_B_*H and MEA*_B_*S).

| **Species** | **DEC (‰)** | **RPP**  **p-value** | **Host MEA_B_ (‰^2^)** | **Symbiont MEA_B_ (‰^2^)** | **Area of overlap (‰^2^)** | **MEA_B_H** | **MEA_B_S** |
| --- | --- | --- | --- | --- | --- | --- | --- |
| *T. gigas* | 0.50 | 0.116 | 6.97 | 2.61 | 2.31 | 0.32 | 0.98 |
| *T. derasa* | 0.93 | <0.001 | 3.65 | 2.02 | 1.63 | 0.41 | 0.82 |
| *H. porcellanus* | 1.14 | <0.001 | 0.78 | 2.15 | 0.41 | 0.49 | 0.17 |
| *T. maxima* | 1.27 | <0.001 | 2.37 | 3.97 | 0.82 | 0.24 | 0.20 |
| *H. hippopus* | 1.57 | <0.001 | 4.05 | 5.28 | 2.01 | 0.43 | 0.34 |
| *T. squamosa* | 1.89 | <0.001 | 5.56 | 5.26 | 1.42 | 0.28 | 0.31 |

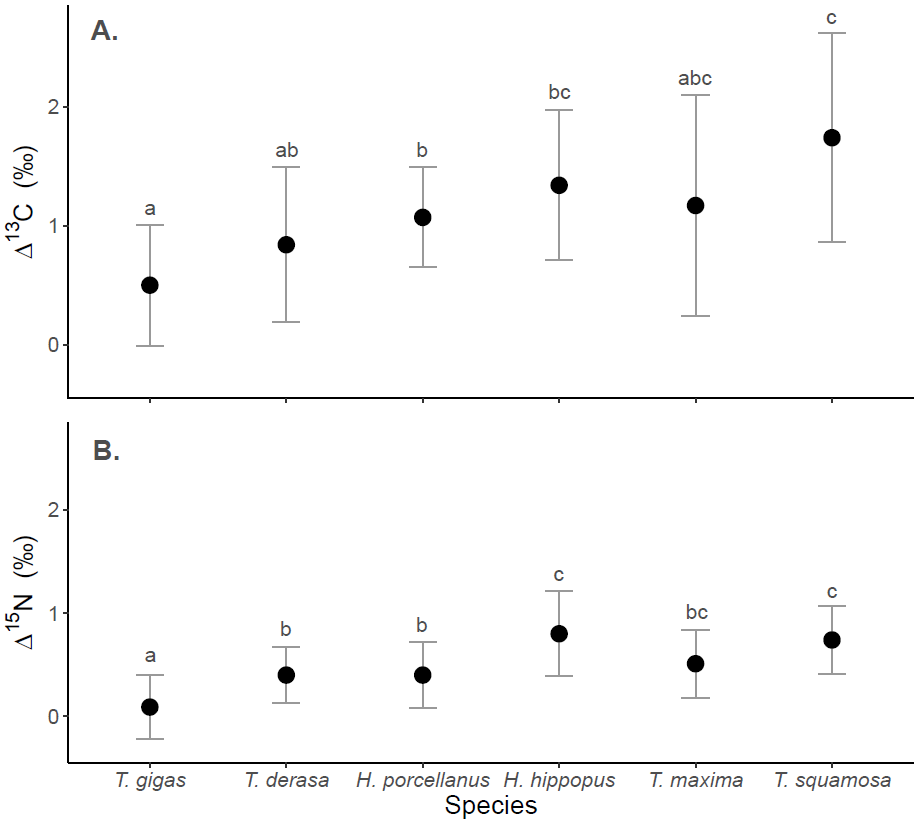

**Supplementary Figure 1.** Mean Δ^13^C (*δ*^13^C_Host_-*δ*^13^C_Symbiont_; **A**) and Δ^15^N (*δ*^15^N_Host_-*δ*^15^N_Symbiont_; **B**) of each giant clam species. Error bars represent one standard deviation. Different letters indicate significant differences between species (ANOVA; *P* < 0.05).

Symbiodiniaceae diversity and community structure

The ITS2 profiles belonging to three Symbiodiniacea genera were found to be different among the six species of giant clams studied (Supplementary Fig. 2). *H. hippopus* and *T. squamosa* associated with four ITS2 type profiles while *T. gigas* and *T. maxima* each hosted three profiles. Both *T. maxima* and *T. squamosa* were dominated by profiles C1/C1c/C3-C1al-C1b (74.95% and 73.45%, respectively) and D4/D5-D9b-D1bq-D5n (17.33% and 18.18%, respectively). Symbiodiniaceae from *T. gigas* predominantly consisted of two different profiles from the genus *Cladocopium* (C93a-C93e-C55a, 62.08% and C93/66-C93e, 31.48%) while *H. hippopus* was mainly composed of profiles from *Cladocopium* (C93a-C93e-C55a, 62.99%) and *Symbiodinium* (A3, 30.76%). Ordination of ITS2 sequences and ITS2 type profiles with NMDS revealed that *T. maxima* and *T. squamosa*, and to a lesser extent in *T. gigas*, *T. derasa, H. hippopus* and *H. porcellanus*, clustered together (Supplementary Fig.3).

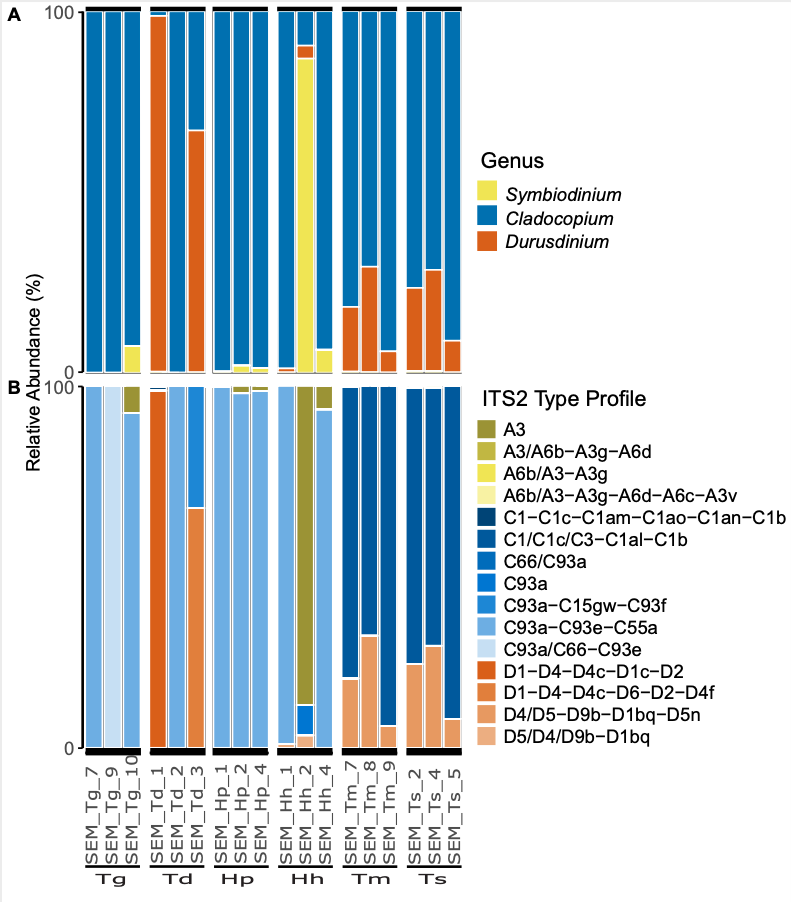

**Supplementary Figure 2. Diversity of Symbiodiniaceae associated with six giant clam species**: *Tridacna gigas* (Tg), *Tridacna derasa* (Td), *Hippopus porcellanus* (Hp), *Hippopus hippopus* (Hh), *Tridacna maxima* (Tm), and *Tridacna squamosa* (Ts). Both relative abundance of Symbiodiniaceae genera (**A**) and predicted ITS2 type profiles (**B**) are shown.

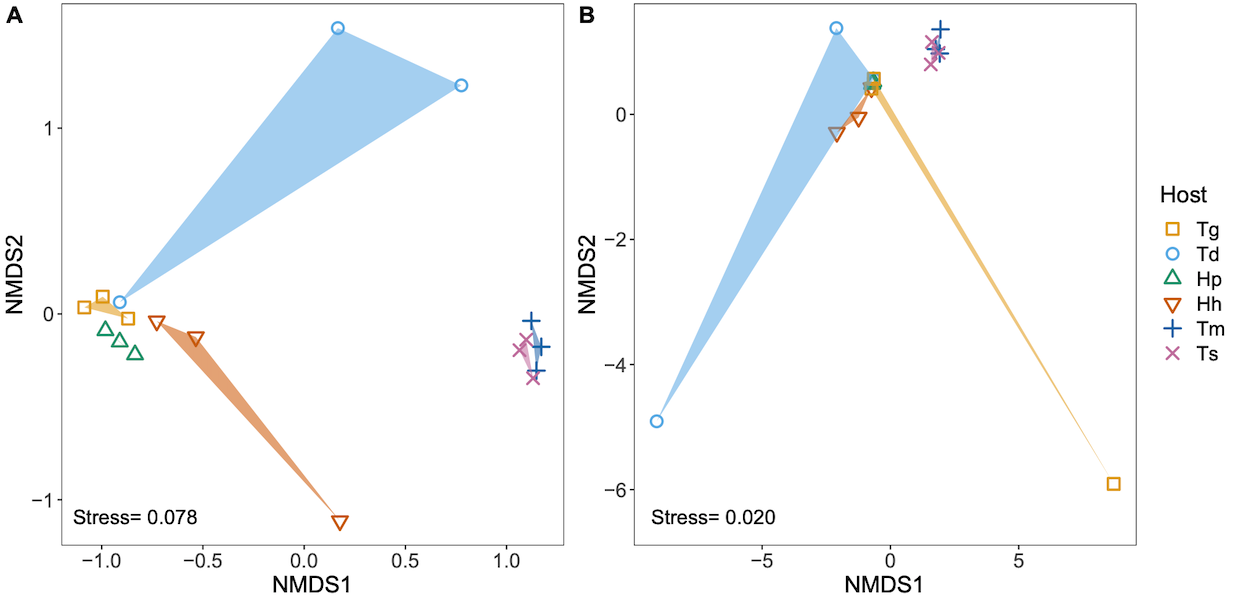

**Supplementary Figure 3**. Non-metric multi-dimensional scaling (NMDS) plots using Bray-Curtis dissimilarities based on (A) ITS2 sequences and (B) ITS2 type profiles of six giant clam species: *Tridacna* *gigas*, *Tridacna* *derasa,* *Hippopus* *porcellanus, Hippopus* *hippopus, Tridacna* *maxima* and *Tridacna* *squamosa*.

**Discussion**

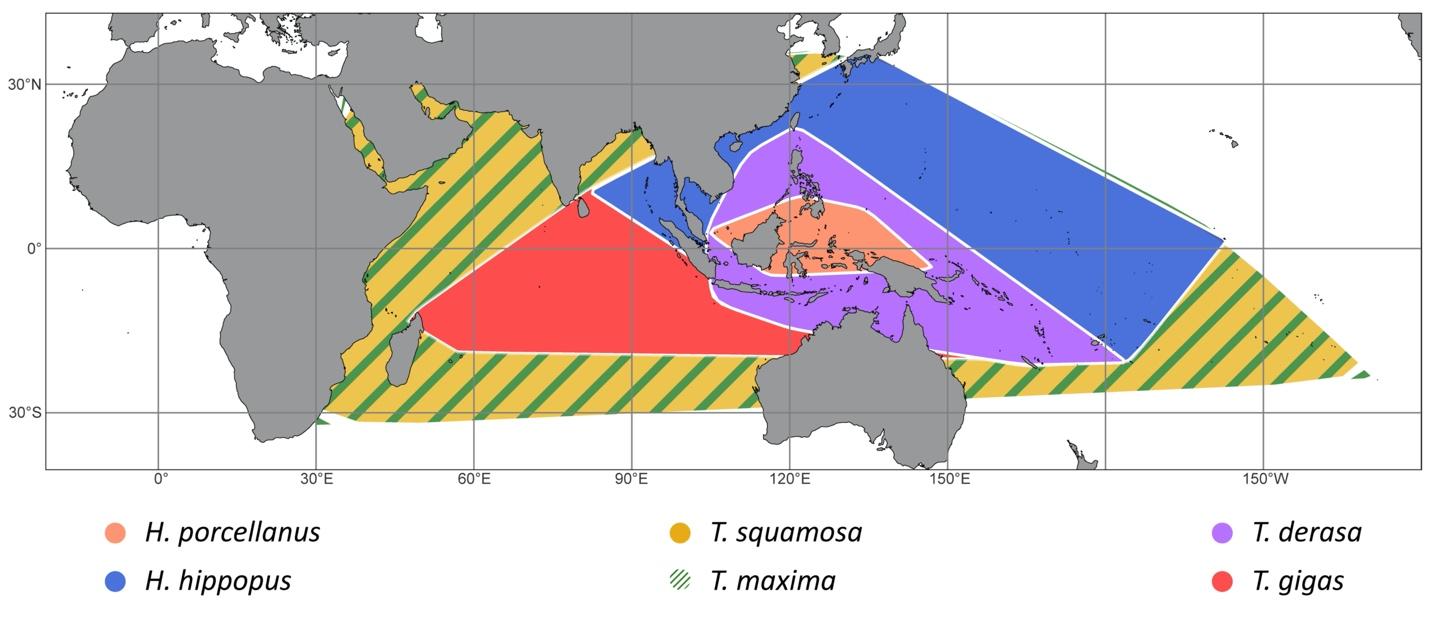

**Supplementary Figure 4.** Geographic distribution of the 6 giant clams species of interest. Orange: *Hippopus porcellanus*, blue: *Hippopus hippopus*, Yellow: *Tridacna squamosa*, Green: *Tridacna maxima*, Purple: *Tridacna derasa*, Red: *Tridana gigas*. Location data were obtained from Neo et al.^1^.

**Materials and Methods**

Sample collection

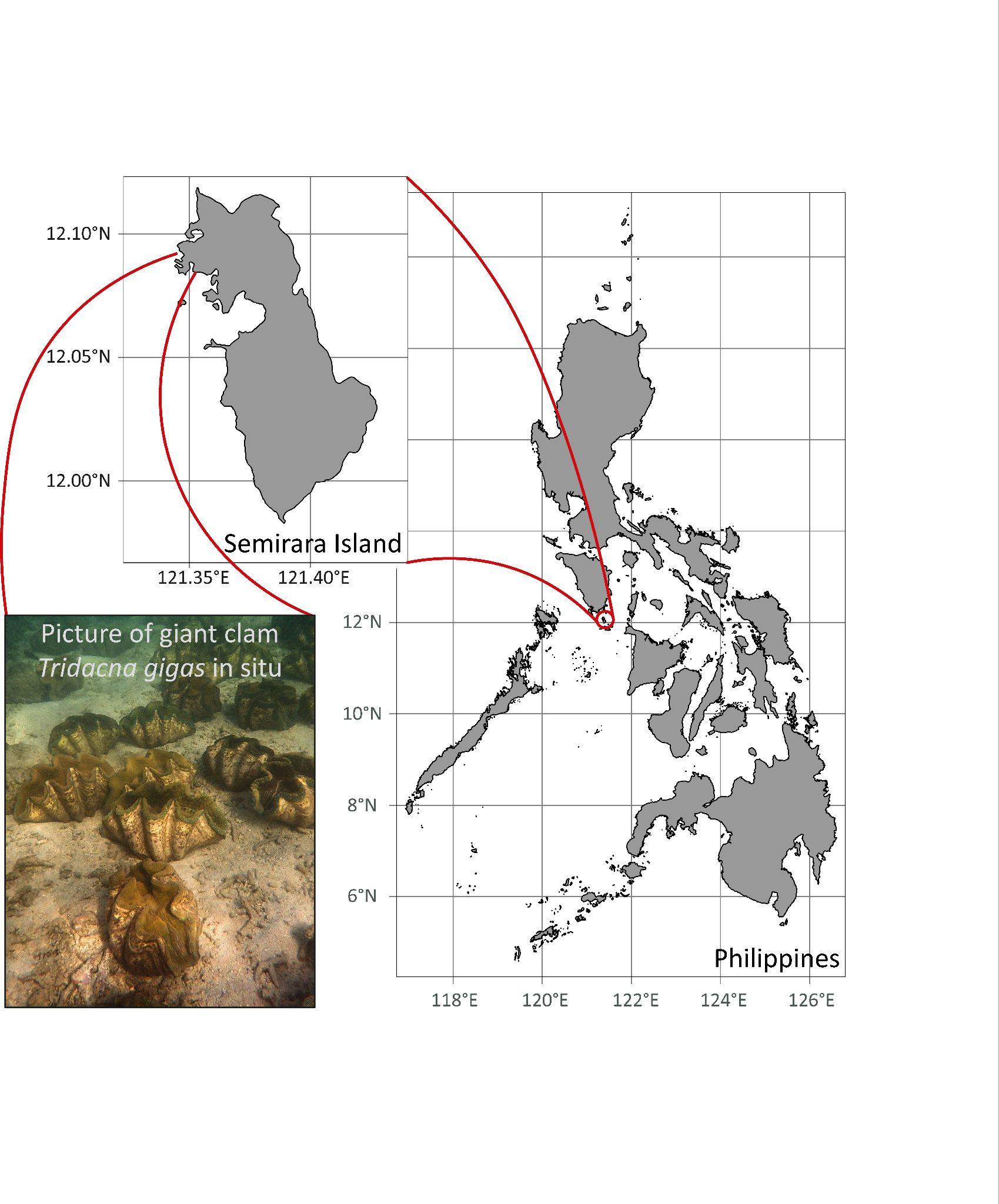

**Supplementary Figure 5.** Map of the study site.

The Semirara hatchery is surrounded by the Tabunan lagoon. Giant clam cohorts were raised at the hatchery in open-circuit raceways with coarse filtered seawater pumped from the Tabunan lagoon before being transferred to sandy areas of the lagoon at 2-5 m depth (Supplementary Fig. 5). Five of the six giant clam species (*Tridacna gigas*, *Tridacna derasa*, *Hippopus porcellanus*, *Hippopus hippopus* and *Tridacna squamosa*) were sampled in the north of the lagoon while the *Tridacna maxima* samples were collected less than 400 m away in the south west of the lagoon. The samples from the five species in the north were collected from a 30 m^2^ area and the *Tridacna maxima* from a 10 m^2^ area. To ensure that both areas had similar conditions, light and temperature hobo loggers were deployed in November 2019. All the samples for SIA analysis were collected across three days in November 2019. On each sampling day, duplicate samples of 3 L of seawater from the Tabunan lagoon were collected and filtered through two 47 mm glass fiber filters (GFF pore size 0.45 μM to collect particulate organic matter. All the filters were placed in a petri dish and dried at ambient temperature before analysis.

Habitat assessment

Light intensity and temperature were highly similar in both areas of the Tabunan lagoon (Supplementary Fig. 6). Visual observations showed that both areas were characterized by sandy sediment, 2-5 m depth and patches of seagrass around the sampled areas.

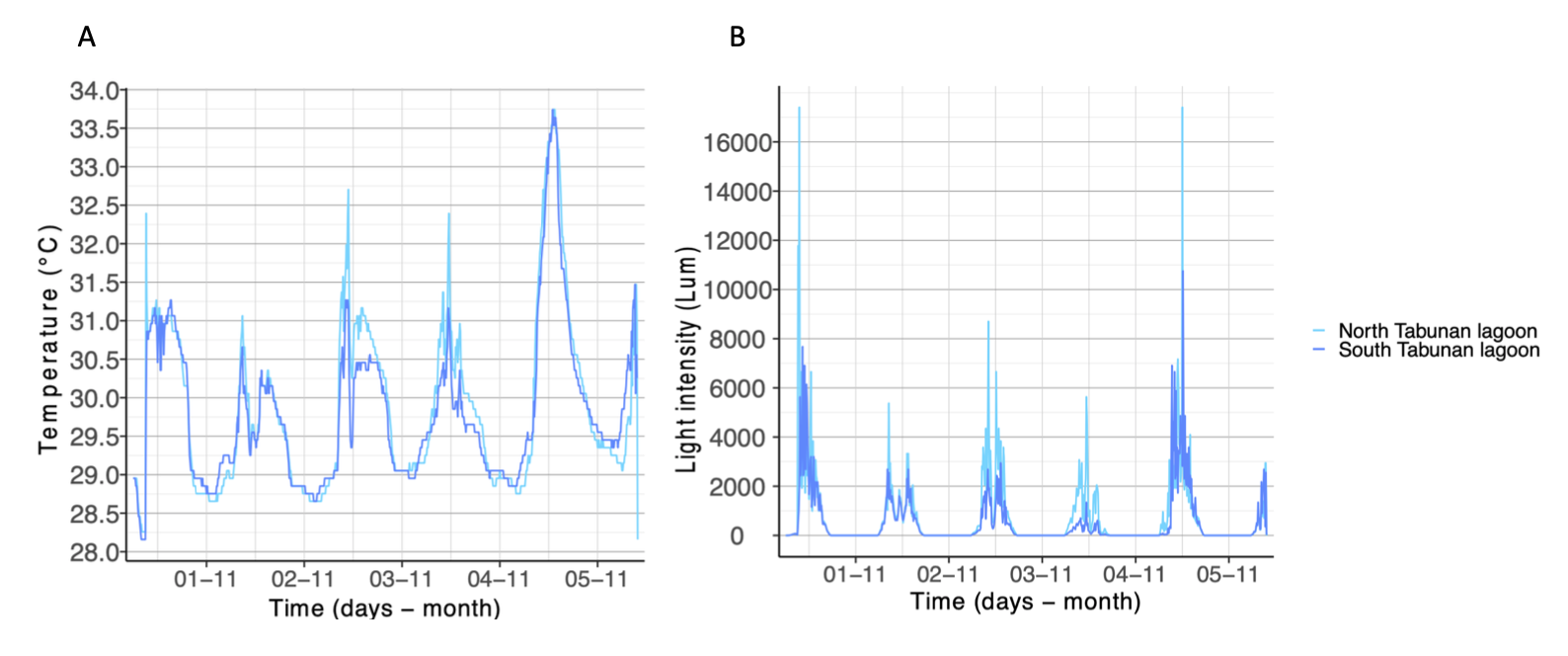

**Supplementary Figure 6.** Light intensity (A) and temperature (B) of the two areas sampled in the Tabunan Lagoon in November.

Laboratory analysis

Amplicon libraries were prepared using a Herculase II Fusion DNA Polymerase Nextera XT Index Kit V2 and sequenced on the Illumina MiSeq platform with 2 x 300 bp paired-end version 3 chemistry. The ITS2 region of the ribosomal DNA operon was used for Symbiodiniaceae community analysis using the primers ITSintfor2^2^ and ITS2-reverse^3^. Triplicate PCR amplifications were carried out under the following conditions: 2 min at 95°C, followed by 35 cycles of 95°C for 30 s, 52°C for 30 s and 72°C for 30 s and a final extension step at 72°C for 5 min^2,3^.

SIA statistical analysis

All statistical analyses and visualizations were executed in R version 4.0.3^4^ using RStudio v.1.4.1103^5^. Stable Isotope Bayesian Ellipses in R (SIBER) analysis was used to fit ellipse areas for sample size (EA) to host and symbiont δ^13^C and δ^15^N values shown on an isotopic biplot(*51*). EAs provide stable results when the number of samples is ≥20(*30*), and thus are used to represent isotopic niche, a proxy for trophic niche. We used a Bayesian analysis to generate a posterior distribution of EA. We used the Markov chain Monte Carlo (MCMC) methods to generate two chains of 20,000 iterations of probable EAs, starting from a vague Inverse Wishart prior and thinned every 10 iterations. The mode of the last 100 posterior draws of EAs was used as the Bayesian estimate of ellipse area (EA_B_)(*60*). We assessed whether the size of clam host isotopic niches differed by comparing the 95% credible intervals of posterior draws of standard ellipse areas (SEA); EAs scaled to fit ±1SD around the mean of each isotope, encompassing 40% of the data(*61*). Species for which the 95% credible intervals did not overlap were deemed to have significantly smaller/larger isotopic niches. Isotopic biplots are presented with EAs derived from the raw data to visually represent results.

Due to discrepancies and omissions in the application of SIBER in the literature, we developed a novel metric that incorporates multiple SIBER outputs to generate one value that quantifies the relative trophic strategy of mixotrophic holobionts. For example, when comparing the location of two groups within isotopic space, different sizes of ellipses have been used with little or no justification. In the initial description of SIBER, Jackson et al. (2011) used SEA (defined above), however their primary goal was to compare the relative sizes of ellipses rather than overlap between ellipses. Subsequent papers have argued for the use of larger ellipses that encompass 95% of the variation of the data (Swanson et al. 2011) and ±2.6 SD around the mean of each variable (major ellipse area; MEA). To conduct the most conservative assessment of nutrient sharing between host and symbionts, we examined the overlap of host and symbiont isotopic niches using Bayesian estimates of used two differently sized ellipses that represent the minimum and maximum of published values: previously used in the literature (Jackson et al 2011; Swanson et al 2015; Skinner et al 2019) to measure the overlap of host and symbiont isotopic niches: (i) SEA_B_s (see above for definition) and (ii) major ellipse area (MEA_B_s) where EA_B_s were scaled to fit ±2.6 SD around the mean of each variable encompassing 95% of data. Employing both ellipse sizes harnesses the bias inherent in each; smaller ellipses are less likely to overlap and thus less likely to detect autotrophy whereas large ellipses are more likely to overlap and less likely to detect heterotrophy. Further, incorporation of both ellipse sizes allows for more fine-scale distinction across species. For instance, a species with overlap between host and symbiont MEA_B_s but not SEA_B_s will be relatively more autotrophic when compared to a species without any overlap between either sets of ellipses. Since both sized ellipses can reveal important information we weighed both equally in our metric. As a first step in quantifying nutrient sharing between symbiotic partners, weTo estimate the contribution of Symbiodiniaceae to the nutrition of each clam host species, determined the area of overlap between host and symbiont niches was determined with the mode of the last 100 posterior draws of EA_B_s.

The second consideration we made when constructing our novel metric was how to standardize the area of overlap across species. Previous work applying SIBER to mixotrophic marine invertebrates expressed the overlap between host and symbiont ellipses as a proportion of host niche area (EA_B_H), representing the contribution symbionts make to host nutrition (Conti-Jerpe et al. 2020; Santos et al. 2021). We chose to also include overlap as a proportion of symbiont niche (EA_B_S) which characterizes the proportion of symbiont photosynthates that are passed to the host. Bayesian overlap estimates were standardized as a proportion of both host SEA_B_ and MEA_B_ metrics (SEA_B_H and MEA_B_H, respectively) and symbiont (SEA_B_ and MEA_B_) ellipses areas. We chose to include both the overlapping proportion of host and symbiont ellipses given the biological relevance of both metrics; EA_B_H represents the contribution of symbiont photosynthesis to overall host trophic niche (proxied by isotopic niche) whereas EA_B_S represents the proportion of symbiont trophic niche that is passed on to the host While symbiont contribution to host niche is a more direct indication of relative host reliance on symbionts, the proportion of symbiont niche that is translocated to the host is indicative of the robustness of the relationship. For example, if a host typically assimilates all the products of their associated symbionts, any decline in symbiont population or productivity will directly impact the daily nutrition of the host and can only be compensated through increasing heterotrophy. Alternatively, a host that uses only a portion of symbiont output could be less impacted and continue to receive the same amount of nutrition during the initial phase of reduced productivity. This thought experiment suggests that EA_B_S should have an exponential impact on the metric score while reaching its maximum. However, relative to EA_B_H, this impact should be mitigated when both overlap values are high since the dependability of the host on its symbionts is secondary compared to the contribution of symbionts to the host nutrition. Similarly, we used the percentage of symbiont SEA_B_ and MEA_B_ overlapping that of the host as an estimate of the proportion of symbiont-produced nutrients assimilated by the host (SEA_B_S and MEA_B_S, respectively). These considerations metrics were incorporated combined into a novel index estimating the relative nutritional importance of the symbionts to the host nutrition called Host Evaluation: Reliance on Symbionts (HERS). In order to increase the stability of the SIBER results obtained, 2000 multiples independant SIBER were computed. The HERS scores were calculated for each analysis and averaged by species. Depending on data used, we recommend to run a minimum of 250 independant SIBER analyses to reach a stable result.

**Data S2. (separate file)**

Stable isotopes data and HERS scores

**Data S3. (separate file)**

Description of the giant clam samples

**Data S4. (separate file)**

Most abundant ITS2 sequences

**Data S5. (separate file)**

Traits for phylogenetic analysis
